# VEGFR3 modulates brain microvessel branching in a mouse model of 22q11.2 deletion syndrome

**DOI:** 10.1101/2022.03.16.484566

**Authors:** Sara Cioffi, Gemma Flore, Stefania Martucciello, Marchesa Bilio, Maria Giuseppina Turturo, Elizabeth Illingworth

## Abstract

The loss of a single copy of *TBX1* accounts for most of the clinical signs and symptoms of 22q11.2 deletion syndrome (22q11.2DS), a common genetic disorder that is characterized by multiple congenital anomalies and brain-related clinical problems, some of which likely have vascular origins. *Tbx1* mutant mice have brain vascular anomalies, thus making them a useful model to gain insights into the brain disorders associated with the human disease. Here, we found that *Tbx1* has a dynamic expression pattern in brain endothelial cells (ECs), including tip cells, during early vascularization, but it is not expressed in EC progenitors. Its main morphogenetic function in the brain is to regulate negatively filopodia biogenesis and vessel branching. Because of similar phenotypes reported for *Vegfr3* loss of function, we pursued a mouse genetic approach to test TBX1-VEGFR3 interaction through gain and loss of function experiments. *Vegfr3* is expressed in brain ECs with extensive overlap with *Tbx1* expression. We demonstrate that inactivating *Vegfr3* in the *Tbx1* expression domain in a *Tbx1* mutant background enhances vessel branching and filopodia formation to a greater extent than that observed in the individual mutants. Furthermore, using a mouse transgenic line, we show that increasing *Vegfr3* expression in the *Tbx1* expression domain fully rescued the vessel branching and filopodia phenotypes caused by *Tbx1* loss of function. Similar results were obtained using an *in vitro* model of endothelial tubulogenesis. Overall, these results provide genetic evidence that *Vegfr3* is a regulator of early vessel branching and filopodia formation in the brain, and is a likely critical effector of the brain vascular phenotype caused by *Tbx1* loss of function.

## INTRODUCTION

Seminal studies in rodents have described the process by which the brain is vascularized through angiogenic sprouting from the perineural vascular plexus. In the embryonic mouse brain, vascularization initiates in the hindbrain around E10 and procedes in a caudal-rostral direction. Within the brain parenchyma, the number of blood vessels increases rapidly to form a dense vascular network that is able to support neurogenesis, which in the mouse initiates around E11.5.

*Tbx1* mutant mice are a model of 22q11.2 deletion syndrome (22q11.2DS), a relativley common chromosome microdeletion disorder, for which most of the clinical problems result from *TBX1* haploinsufficiency (Ogata et al., 2014; Paylor et al., 2006; Torres-Juan et al., 2007; Yagi et al., 2003). In practical terms, the most vexing aspect of the clinical phenotype is the brain-related clinical problems, some of which might have vascular origins. TBX1 is a member of the family of T-box trascription factors that is widely expressed in the embryonic pharynx, and it is also expressed in the endothelial lining of a subset of brain vessels in pre-term mouse embryos (Paylor et al., 2006). We have previously shown that *Tbx1* plays a critical cell-autonomous role in brain endothelial cells (ECs) that determines the correct density and functionality of brain vessels (Cioffi et al., 2014). We have proposed that TBX1 exerts these effects through interactions, direct or indirect, with *Vegfr3* and *Dll4* in ECs, both of which have anti-angiogenic functions in the mouse brain (Suchting et al., 2007; Tammela et al., 2011) and are regulated by TBX1 in ECs, including brain ECs, *in vivo* and *in vitro* (Chen et al., 2010; Cioffi et al., 2014). Endothelial-specific inactivation of each of these genes causes brain vessel hyperbranching (Cioffi et al., 2014; Suchting et al., 2007; Tammela et al., 2011). We have previously shown that the Notch agonist JAG1 is not sufficient to rescue fully endothelial microtubule hyperbranching in cultured ECs silenced for *TBX1* (Cioffi et al., 2014), suggesting that *Vegfr3* and/or Notch-independent pathways are involved.

In this study, we investigated the TBX1-VEGF-C/VEGFR3 axis. We demonstrate that *Tbx1* expression is activated in vascular endothelial cells, rather than in their mesodermal precursors, during early brain vascularization. Expression initiates in the hindbrain neuroepithelium at E10.5, and then spreads to most of the brain by E15.5. We found a broad overlap between TBX1 and VEGFR3 in brain vessels and, importantly, genetic experiments determined that *Tbx1*-driven enhanced expression of *Vegfr3* is sufficient to rescue the brain vasculature phenotype of *Tbx1* mutants. Overall, our results indicate that Vegfr3 is a key effector of the modulatory function of TBX1 in brain vessel branching.

## MATERIALS AND METHODS

### Mouse lines and genotyping

Mouse studies were performed according to the animal protocol 257/2015-PR reviewed by the local IACUC committee and by the Italian Istituto Superiore di Sanità and approved by the Italian Ministero della Salute, according to Italian law and European guidelines. The following mouse lines were used: *Tbx1*^*lacZ/+*^ (Lindsay et al., 2001), *Tbx1*^*Cre/+*^ (Huynh et al., 2007), *Vegfr3*^*flox/+*^ (Zarkada et al., 2015), *Vegfr3*^*+/-*^ (Martucciello et al., 2020), *Rosa*^*mTmG*^ (Muzumdar et al., 2007), TgVegfr3 (Martucciello et al., 2020). Genotyping of mice was performed as in the original reports.

### Immunofluorescence on brain sections

#### Cryosections

E9.5, E10.5, E11.5 embryos were fixed in 4% PFA/1x PBS at 4°C overnight, washed in 1x PBS and incubated for 12 h in serial dilutions of sucrose/1x PBS (10%, 20%, 30% sucrose) at 4°C. Brains were then incubated for 2 h at 4°C in 50:50 v/v 30% sucrose/1x PBS/OCT prior to embedding in OCT. Ten micron coronal sections were cut along the rostral–caudal brain axis on a cryotome. Alternatively, specimens were stored at −80°C. Experiments were performed on serial sections 200 μm apart (5 embryos/genotype). Sections were briefly microwaved to boiling point in 10 mM sodium citrate (pH 6.0) 3 times, for antigen enhancement, cooled, rinsed in 1x PBS, permeabilized with 0.1% Triton x-100 blocked in 10% goat serum (GS) /1x PBS /0.1% Triton x-100 for 1h at RT.

#### Thick brain sections

E13.5, E15.5, E18.5 mouse brains were fixed in 4% PFA at 4°C overnight and subsequently embedded in 4% low melting agarose in 1x PBS. Serial thick coronal sections of 50 μm (E13.5) or 100 μm (E15.5, E18.5) were cut on a vibratome at 4°C. Brains (fixed) that were not used immediately were stored at 4°C in 0.0025% NaNO3.

Immunofluorescence was performed on serial sections along the rostral–caudal brain axis using the following antibodies: mouse monoclonal anti-GLUT1 (Abcam, ab40084), chicken polyclonal anti-GFP (Abcam, ab13970,), anti-mouse monoclonal VEGFR3/FLT4 (eBioscience, clone AFL4, Cat. #14-5988-85), rabbit polyclonal anti-VEGFR3/FLT4 (Elabscience, #36398), rat anti-mouse monoclonal VEGFR2/FLK1 (BD Pharmingen, Avas 12α1, Cat. #550549). Secondary antibodies used were goat anti-mouse Alexa Fluor 594 and 488, goat anti-rabbit Alexa Fluor 488 and 594, goat anti-rat Alexa Fluor 594, goat anti-chicken Alexa Fluor 488. Sections were then incubated with primary antibodies overnight at 4°C in the same blocking solution reducing the GS to 5%, rinsed, and incubated in secondary antibodies for 1h 30 min at RT. Fluorescence was observed with an epifluorescence microscope (Leica DMI6000B, acquisition software LAS AF 2.6 or Nikon Confocal Microscope A1, mounted on Nikon ECLIPSE Ti, acquisition software NiS element. Images were digitally documented with a camera and computer processed using Adobe Photoshop® version 6 for Windows.

### Quantitative analysis of vascular anomalies

All quantitative analyses were performed on three brain sections per embryo in rostral, medial and caudal postions that correspond visually to the following stereotaxic levels (Paxinos G, and Franklin K., 2012). Rostral: Bregma 0.74 mm; Medial: Bregma -2.54 mm; Caudal: Bregma -4.84 mm, where R: encompasses the dorsal and ventral sub-pallium; M: the dorsal and ventral thalamus; C: the dorsal and ventral midbrain.

### Vessel branch point and filopodia counts

We analyzed a minimum of 5 embryos per genotype in all experiments. Flattened images were digitally reconstructed from confocal *z* stacks of 2.5 μm representing three different brain regions (rostral, medial and caudal). For each flattened image, we manually counted all the branch points in a single field per section, and thus three fields per embryo. The area counted per field was 1.444 mm^2^ and the total length of the vascular network per field was measured. For filopodia, we counted all the filopodia in two sub-fields per section (rostral, medial and caudal) for a total of six fields per embryo. The area counted per field was 0.045 mm^2^ and counts are expressed as the mean number of filopodia per 100 μm of vessel length. Counts were performed using the ImageJ cell counter plugin.

### Statistical analysis

The statistical analysis of the data pertaining to the brain vessel density was performed using a likelihood ratio test for Negative Binomial generalized linear models. We first calculated the mean number of branchpoints / field / embryo. We then calculated the mean for the group (embryos / genotype). The latter value was used for the statistical analysis. For the analysis of data pertaining to filopodia we performed the Kruskal Wallis test followed by the unpaired Mann Whitney U test.

### Cell manipulations

HUVECs (Lonza) were transfected by Lipofectamine™ 2000 (Invitrogen). For *TBX1* or *VEGFR3* knockdown (kd) in HUVECs, RNA interference was performed using a commercial siRNA for *TBX1* or *VEGFR3* (ON-TARGETplus SMARTpool, Thermo Fisher Scientific) (40 nM) or a control (non-targeted) siRNA (Thermo Fisher Scientific) as previously described (Cioffi et al., 2014). mRNA expression was evaluated by qReal Time PCR. For *Vegfr3* overexpression, HUVECs were transfected with 1μg of pcDNA3 plasmid vector containing a full-length mouse *Vegfr3* cDNA (GeneCopoeia), or with empty vector (Martucciello et al., 2020). Protein expression was evaluated by western blotting with anti-VEGFR3 antibody. Twentyfour hours after transfection, cells were collected and processed for matrigel assay.

For *Vegfr3* over-expression in *Tbx1*^*KD*^ endothelial cells, 24 h after transfection of HUVECs with a *TBX1* or control siRNAs, cells were transfected with pcDNA3-*Vegfr3*. Twentyfour hours after the second transfection, cells were collected and processed for matrigel assays. *Tbx1* and *Vegfr3* mRNA expression was evaluated by qReal Time PCR.

### Matrigel assays

200 ml of Matrigel (BD Bioscience) was plated onto chilled 15-mm wells and incubated at 37°C for 30 min, as per the manufacturer’s instructions. HUVECs in 6-well plates, previously transfected as described above, were trypsinized and counted. 1.5 × 10^5^ treated cells in EGM-2 media (Lonza) were added to each well containing Matrigel. After 16 h at 37°C, the formation of microtubules was analyzed using an Olympus CKX41 Image Analyzer. The quantification of microtubule branch points was performed after dividing each large image into nine sub-images. The number of branch points was calculated as the sum of counts made in all nine sub-images. Quantitative analysis was performed using the Image J software.

## RESULTS

### Tbx1 expression is activated in ECs rather than in EC progenitors

Many of the TBX1-dependent functions in the cardiovascular system are linked to progenitors rather than to differentiated cells. As *Tbx1* is widely expressed in the head mesoderm, we addressed the question of whether the gene is activated in EC precursors before the neuroepithelium is vascularized. To this end, we crossed *Tbx1*^*Cre/+*^ mice with *Rosa*^*mTmG*^ reporter mice (Muzumdar et al., 2007) and evaluated the distribution of Tbx1-fated cells in serial coronal brain sections of *Tbx1*^*Cre/+;*^ *Rosa*^*mTmG*^ embryos between stages E9.5 and E18.5, where these cells are marked by *Tbx1*^*Cre*^-activated expression of GFP. Sections were co-immunostained with anti-KDR (E9.5 - E11.5) or anti-Glut1 (E13.5 - E18.5) to identify ECs. Overall, results showed that *Tbx1*-expressing cells and their progeny populate the developing brain following caudal to rostral and ventral to dorsal trajectories, as shown in the cartoon in Fig. 2M. Specifically, at E9.5, a few ECs (KDR+) in the perineural vascular plexus (PNVP) were GFP+, while the surrounding head mesenchyme was heavily populated with GFP+ cells (Fig. 1A). The first GFP+ cells localized to blood vessels (KDR+) within the hindbrain neuroepithelium at E10-E10.5 (Fig. 1B). Between E10.5 and E11.5, the number of GFP+ cells increased dramatically in the hindbrain neuroepithelium, such that by E11.5, co-expression of GFP and KDR was almost 100% (Fig. 1C). However, at this stage, the midbrain (Fig. 1D) and forebrain (Fig. 1E) contained only a few isolated GFP+;KDR+ cells. At E13.5 (Fig. 2A-F), the presence GFP+;GLUT1+ cells increased in the ventral forebrain (Fig. 2ab’, 2cd’) and ventral midbrain (Fig. 2ef’), but the dorsal forebrain (Fig. 2ab, 2cd) and dorsal midbrain (Fig. 2ef) remained largely free of GFP+ cells. The contribution to and distribution of GFP+ cells in the aforementioned brain structures were maintained at E15.5 (Fig. 2G-L) in the forebrain (boxed areas in Fig. 2G and 2I) and in the midbrain (boxed area Fig. 2K), and at E18.5 (Supplementary Fig. S1, A-F), suggesting that *Tbx1* is not activated significantly in dorsal brain regions during embryonic development (Supplementary Fig. S1, A, C, E). At no developmental stage were GFP+/KDR-negative or GFP+/GLUT1-negative cells seen within the brain parenchyma, confirming that Tbx1-fated cells are endothelial cells.

**Figure 1.**
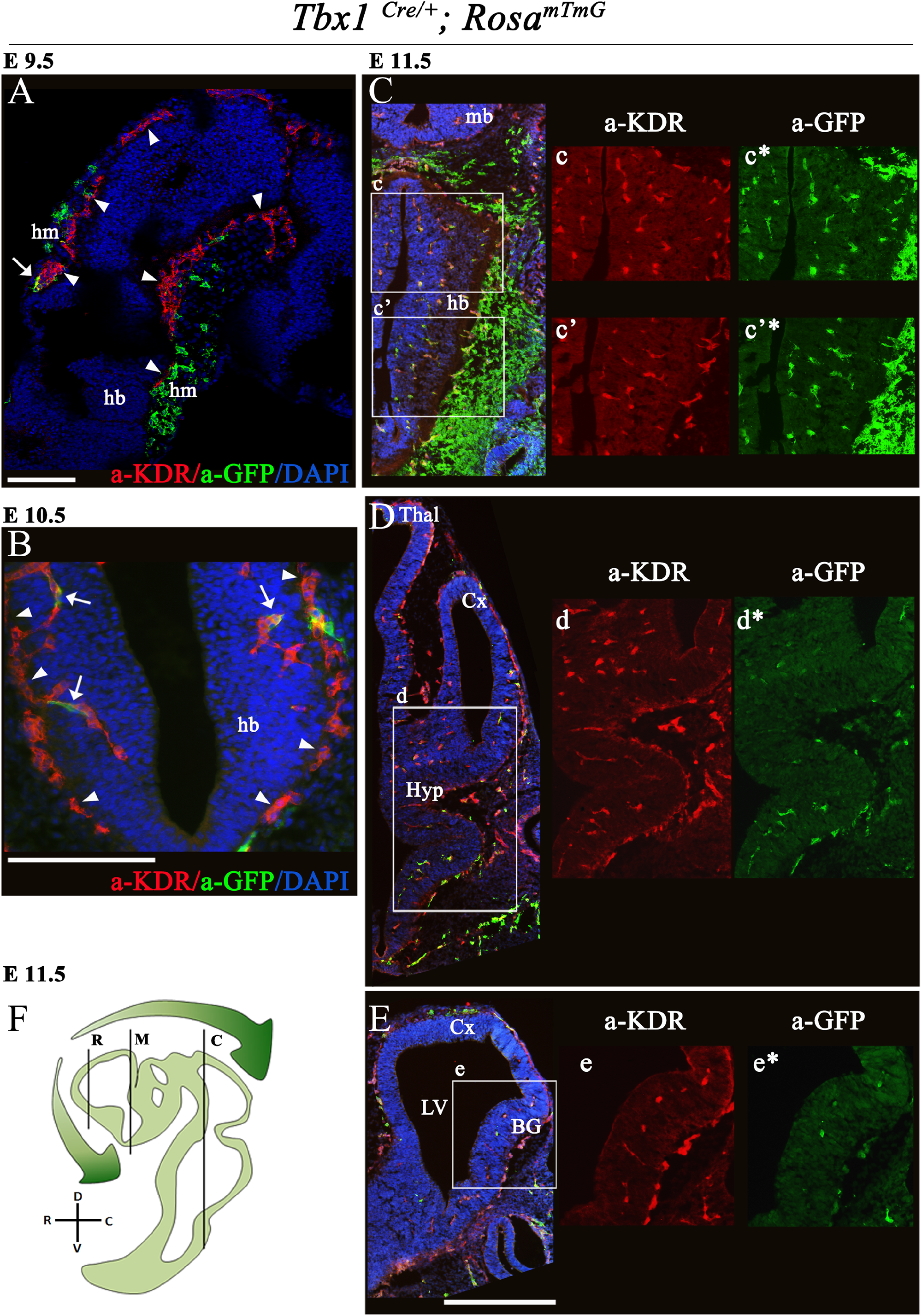
Distribution of Tbx1-fated cells in the brain of Tbx1^Cre/+;^ Rosa^mTmG^ embryos. (A - E) Representative coronal brain sections of embryos at E9.5 (A), E10.5 (B) and E11.5 (C, caudal section), (D, medial section), (E, rostral section) immunostained for GFP (green) and KDR (red). White arrowheads in A and B indicate the perineural vascular plexus and white arrows cells co-expressing GFP and KDR. White boxes in C-E (merge) are shown as enlarged, single color channels to the right. (F) the cartoon shows the position of the coronal brain sections shown in panels C (caudal, C), D (medial, M) and E (rostral, R). Green shaded arrows indicate the relative density of GFP+ cells (*Tbx1*-fated) along the rostral --> caudal brain axis at E11.5. Scale bars, A, 250 μm, B-E, 500 μm Abbreviations: hm, head mesenchyme, hb, hindbrain, Cx, cortex, LV, lateral ventricle, BG, basal ganglia, mb, midbrain, Thal, thalamus, Hyp, hypothalamus.

**Figure 2.**
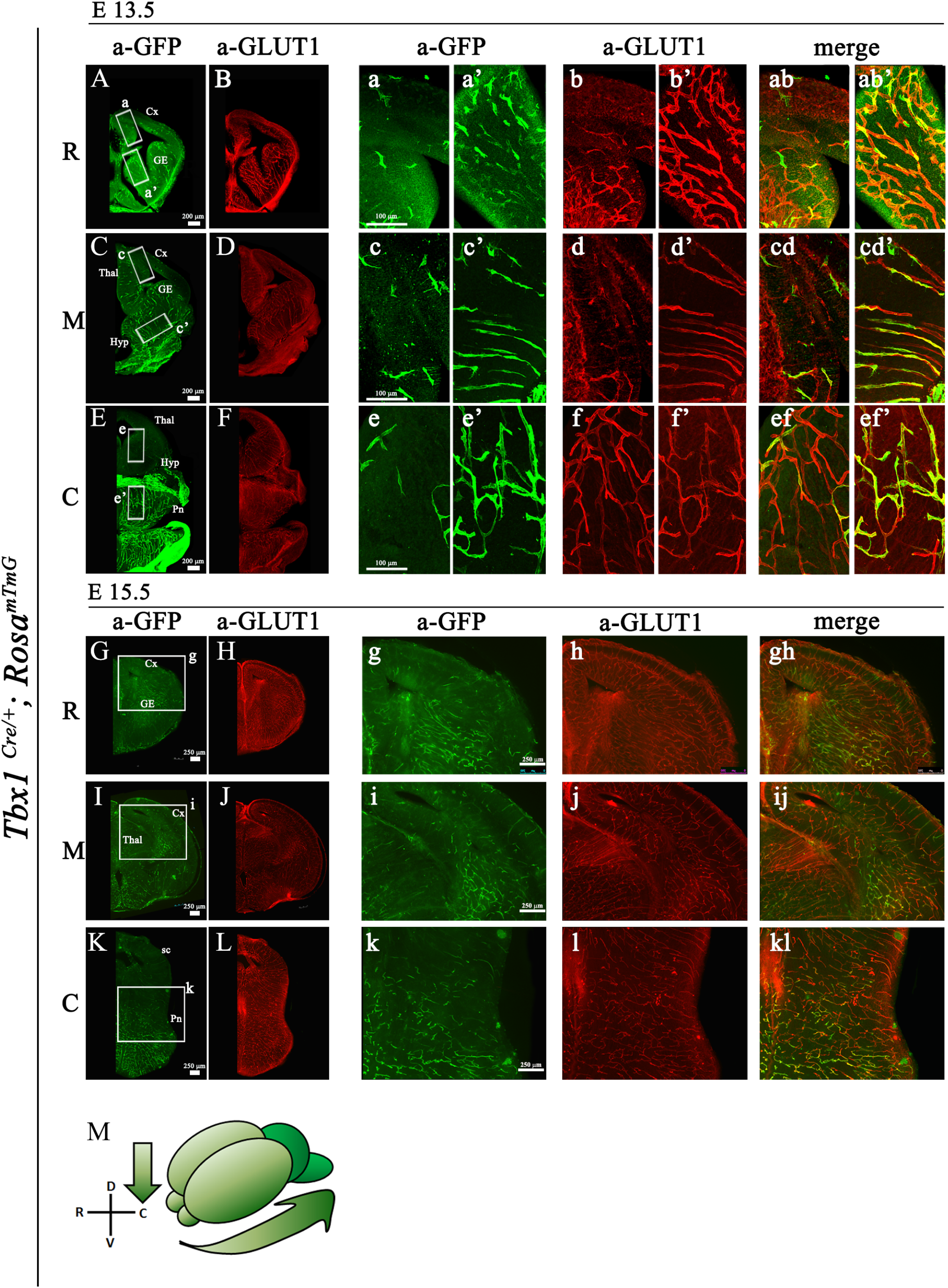
Distribution of Tbx1-fated cells in the brain of Tbx1^Cre/+;^ Rosa^mTmG^ embryos. Representative coronal brain sections of embryos at E13.5 (A - F) and E15.5 (G - L) immunostained for GFP (green) and GLUT1 (red). Boxed areas in A, C, E, G, I, K are enlarged in the adjacent panels. (M) cartoon showing the relative density of GFP+ cells along the rostral --> caudal and dorsal --> ventral axes. Abbreviations: R, rostral, M, medial, C, caudal, Cx, cortex, GE, ganglionic eminences, Thal, thalamus, Hyp, hypothalamus, Pn, pons, sc, superior colliculus.

### Tbx1 and VEGFR3 co-localize in brain ECs, including tip cells

In order to determine the extent of overlap between TBX1 and VEGFR3, we first determined whether the anti-VEGFR3 antibody used in this study showed pan-endothelial expression in the mouse brain as reported (Watanabe et al., 2019). For this, and for the subsequent analysis of TBX1-GFP and VEGFR3 co-expression, we used the same sectional series of *Tbx1*^*Cre/+;*^ *Rosa*^*mTmG*^ embryos. Co-immunostaining with anti-GLUT1 and anti-VEGFR3 antibodies revealed overlapping expression of the two proteins in virtually all brain vessels along the rostral-caudal brain axis at E13.5 (Supplementary Fig. S1, G-L). We next co-immunostained an adjacent series of sections with anti-GFP and anti-VEGFR3 antibodies. This revealed extensive co-localization of the proteins in brain vessels along the rostral-caudal brain axis at the stages analyzed, namely E13.5 (Fig. 3A-F, 3ab, 3cd, 3ef, 3ef’), E15.5 (Fig. 3G-L, 3gh, 3ij, 3kl) and E18.5 (Supplementary Fig. S1, M - R). We then asked whether the co-localization of TBX1-GFP and VEGFR3 in brain vessels included endothelial tip cells, which are distinguished from stalk cells by the presence of filopodia. Anti-GFP immunostaining of brain sections of *Tbx1*^*Cre/+*^; *Rosa*^*mTmG*^ embryos at E13.5 and E15.5 labelled endothelial tip cells, including their filopodial extensions (Fig. 3M-P). We found that many but not all GFP+ tip cells expressed VEGFR3, but two different anti-VEGFR3 antibodies failed to label the filopodia (Fig. 3N, 3P). This may be due to an inability of the antibodies to bind to the receptor in this territory, or to a lack of the receptor at this location in these mutants.

**Figure 3.**
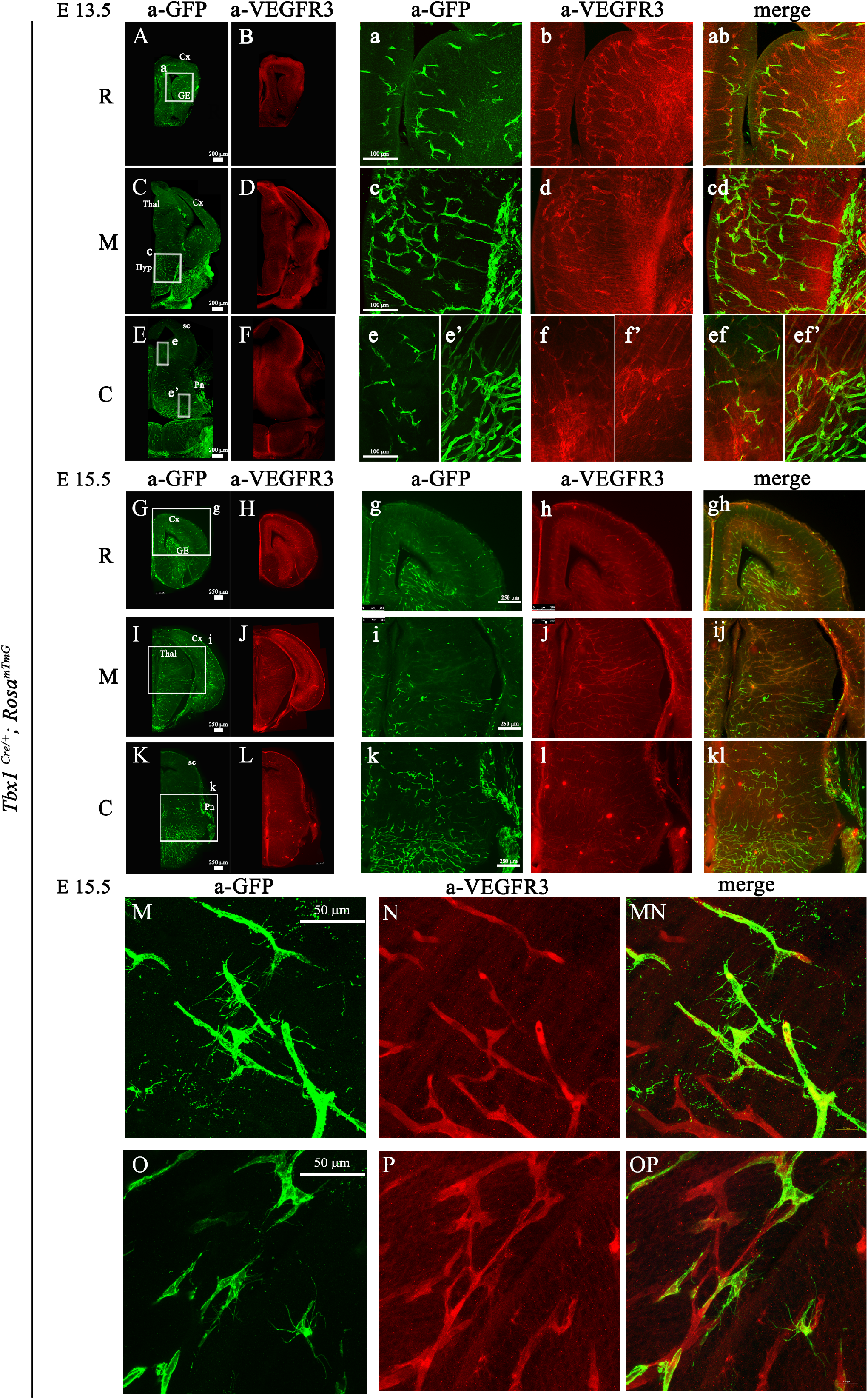
TBX1 and VEGFR3 colocalize in brain endothelial cells. Representative coronal brain sections of *Tbx1*^*Cre/+;*^ *Rosa*^*mTmG*^ embryos at E13.5 (A - F) and E15.5 (G - P) immunostained for TBX1-GFP (green) and VEGFR3 (red). Boxed areas in A, C, E, G, I, K are enlarged in the adjacent panels. (M-P) High magnification images show that endothelial filopodia are labelled by anti-GFP (M, O) but not anti-VEGFR3 (N, P, MN, OP) antibodies. Abbreviations: R, rostral, M, medial, C, caudal, Cx, cortex, GE, ganglionic eminences, Thal, thalamus, Hyp, hypothalamus, Pn, pons, sc, superior colliculus.

### Vegfr3 haploinsufficiency enhances the angiogenic sprouting phenotype of Tbx1 mutants

We next tested whether *Tbx1* and *Vegfr3* interact genetically. For this, we intercrossed *Tbx1*^*Cre/+*^ (*Tbx1*^*Cre*^ is a null allele) and *Vegfr3*^*flox/+*^ mice and quantified brain vessel branchpoint density and angiogenic sprouting (filopodial density) in thick (100 μm) brain cryosections of conditional compound heterozygous embryos at E18.5 immunostained with anti-GLUT1 (Fig. 4A-C, Supplementary Fig. S2). Results revealed an increased density of endothelial filopodia in *Tbx1*^*Cre/+*^;*Vegfr3*^*flox/+*^ embryos compared to *Tbx1*^*Cre/+*^ embryos (Fig. 4B, 4C, 4f, *P* = 0.0317 or WT controls *P* = 0.0079), but there was no difference in branchpoint density (Fig. 4F). Thus, *Vegfr3* heterozygosity enhances the brain vasculature phenotype in *Tbx1* heterozygous (*Tbx1*^*Cre/+*^) mutants, albeit mildly. Interestingly, a similar result was obtained in *Vegfr3*^*+/-*^ (Fig. 4E) embryos (germline heterozygous) versus WT embryos (Fig. 4F’, 4f’), i.e., increased filopodial density (*P* = 0.0079) but unaltered branchpoint density (*P* = 0.117), indicating that there might be non-productive angiogenic sprouting in *Vegfr3*^*+/-*^ embryos. As far as we know, this is the first time that *Vegfr3* has been reported to be haploinsufficient.

**Figure 4.**
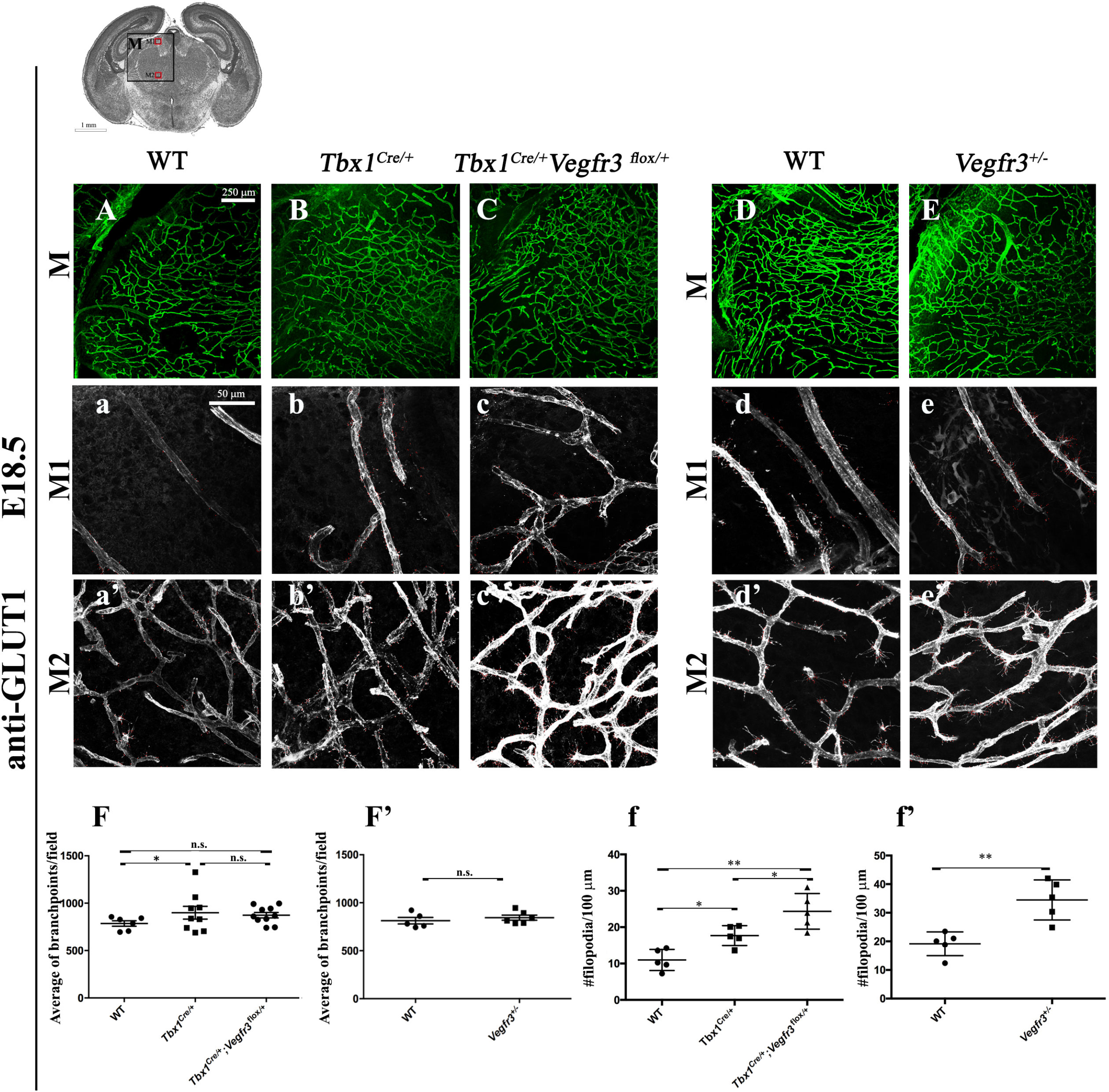
Tbx1 and Vegfr3 interact genetically to regulate brain vessel and filopodial density. (A - E) Representative coronal brain sections (medial) of E18.5 embryos immunostained for GLUT1. The cartoon indicates the position of the counting boxes (M1, M2), shown at high magnification in panels M1 (a - e) and M2 (a’ - e’), used to quantify vessel branchpoint (F, F’) and filopodial (f, f’) density in embryos with the indicated genotypes. *** P value <0.001, ** P value <0.01, * P value <0.05. Error bars ± SD. Abbreviations: Cx, cortex, hp, hippocampus, Thal, thalamus.

The extensive co-localization of Tbx1-GFP and VEGFR3 in brain vessels, together with the predominantly ventral distribution of Tbx1-fated cells (GFP+), suggest that ventral rather than dorsal brain structures may be the preferred sites of TBX1-VEGFR3 interaction, and in particular the ventral hindbrain, where TBX1 is highly expressed. VEGFR3 was expressed in most brain vessels already at E13.5, (this study and (Watanabe et al., 2019)) but, brain vessel density was normal in *Tbx1* homozygous embryos at this stage (*P* = 0.07, not shown), suggesting that the critical interaction between the two genes likely occurs between E13.5 and E15.5.

### Enhanced expression of Vegfr3 in Tbx1-depleted ECs rescues microtubule hyperbranching

Because reduced dosage of *Vegfr3* enhances the *Tbx1* mutant phenotype, we reasoned that VEGFR3 may be an effector of TBX1 in regulating brain vessel density. If this were the case, supplemental *Vegfr3* expression should rescue the phenotypic consequences of *Tbx1* mutation. The likelyhood of success of this approach depends upon the effects of altered *Vegfr3* dosage on EC proliferation and vessel growth, but *in vivo* studies indicate that VEGFR3 has both pro- and anti-angiogenic potential (Karkkainen et al., 2000; Karkkainen et al., 2001; Mäkinen et al., 2001). Therefore, prior to initiating *in vivo* rescue experiments, we first tested whether isolated ECs respond to *Vegfr3* overexpression in a way that would predict *in vivo* rescue. We tested this in a functional assay that contains only ECs (HUVECs) cultured in Matrigel™ (Fig. 5). We have shown previously that *TBX1* knockdown in HUVECs causes microtubule hyperbranching in Matrigel™ endothelial cultures (Cioffi et al., 2014). Thus, results obtained in the cell-based assay were consistent with *in vivo* data obtained in the same study in *Tbx1* mutant mice, which showed brain vessel hyperplasia (Cioffi et al., 2014). In Matrigel assays, we found that knockdown of *Vegfr3* by transfection with a *VEGFR3* targeting siRNA led to an approximately 60% increase in microtubule branching compared to HUVECs transfected with a control siRNA (Fig. 5A, *P* = 0.0138), while over-expression of *Vegfr3* in HUVECs, achieved by transfection with a plasmid containing a full-length murine *Vegfr3* cDNA (pcDNA3-Vegfr3), led to microtubule hypobranching; reduced by approximately 24% compared to HUVECs transfected with the empty plasmid vector (Fig. 5B, *P* = 0.0027). We then tested whether over-expression of *Vegfr3* would rescue microtubule hyperbranching in HUVECs after *TBX1* knockdown. For this, we co-transfected HUVECs with pcDNA3-Vegfr3 and siRNA-TBX1 or siRNA-CTR (control) and measured microtubule branching 12 h after plating in Matrigel™ and 24 h after the second transfection. Results showed that over-expression of *Vegfr3* partially rescued microtubule hyperbranching in HUVECs after *TBX1* knockdown, although it did not restore branching to wild type levels (Fig. 5C, *P* = 0.0148).

**Figure 5.**
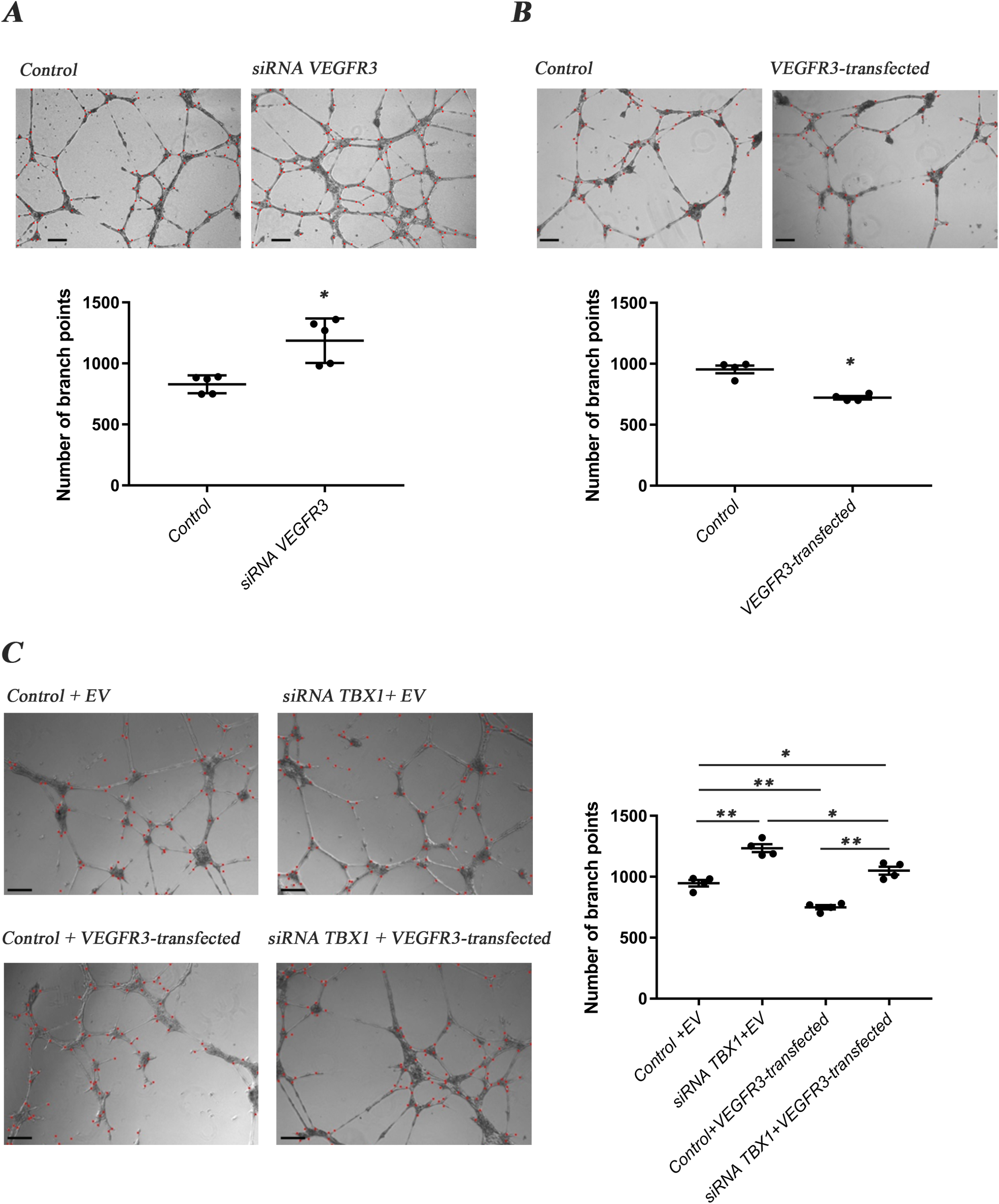
Vegfr3 overexpression represses formation of endothelial microtubules. A) microtubule hypobranching after trasfection with pcDNA3-Vegfr3, B) hyperbranching after VEGFR3-targeted siRNA, C) pcDNA3-Vegfr3 rescues hyperbranching caused by *TBX1* knockdown. *** *P* value <0.001, ** *P* value <0.01, * *P* value <0.05. Error bars ± SD. Scale bar, 200 μm. Abbreviations: EV, empty vector.

### Transgenic expression of Vegfr3 rescues brain vessel hyperplasia caused by loss of Tbx1 function in vivo

We asked whether over-expression of *Vegfr3* in the *Tbx1* expression domain would rescue the brain vessel abnormalities of *Tbx1* mutant embryos. To this end, we crossed *Tbx1*^*Cre/+*^ mice with mice carrying a single copy of a Cre-activatable murine *Vegfr3* transgene, TgVegfr3 mice (Martucciello et al., 2020). We then intercrossed mice with the genotypes *Tbx1*^*Cre/+*^;TgVegfr3 and *Tbx1*^*lacZ/+*^ and measured brain vessel density and filopodial density in E18.5 embryos with the following genotypes: *Tbx1*^*+/+*^, *Tbx1*^*Cre/+*^, *Tbx1*^*Cre/lacZ*^, TgVegfr3;*Tbx1*^*Cre/+*^ and TgVegfr3;*Tbx1*^*Cre/lacZ*^ that were immunostained with anti-GLUT1 (Fig. 6, Supplementary Fig. 3). Results revealed the presence of brain vessel hyperplasia (+17%, *P* = 0.0365) and increased filopodial density (+85%, *P* = 0.0079) in *Tbx1* heterozygous (*Tbx1*^*Cre/+*^) embryos (Fig. 6B - 6b’, 6F, 6f) and in *Tbx1* homozygous embryos (+35.5% branchpoints, *P* = 2.126e-09 and +58% filopodia, *P* = 0.0079 respectively (Fig. 6D - 6d’, 6F, 6f). More importantly, *Tbx1*^*Cre/+*^-induced activation of the *Vegfr3* transgene fully rescued both brain vessel phenotypes in *Tbx1*^*Cre/+*^;TgVegfr3 (heterozygous for *Tbx1*) embryos (Fig. 6C - 6c’) and *Tbx1*^*Cre/lacZ*^;TgVegfr3 (homozygous for *Tbx1*) embryos compared to controls (6E - 6e’, 6F), i.e., they returned to wild type levels. As *Tbx1*^*Cre*^*-*driven recombination in the mouse brain is limited to ECs (shown in Fig. 2), these results indicate that *Tbx1*-driven activation of *Vegfr3* transgene in brain vessels is sufficient to compensate for the loss of function of Tbx1 and rescues the normal brain vessel density in this mouse model.

**Figure 6.**
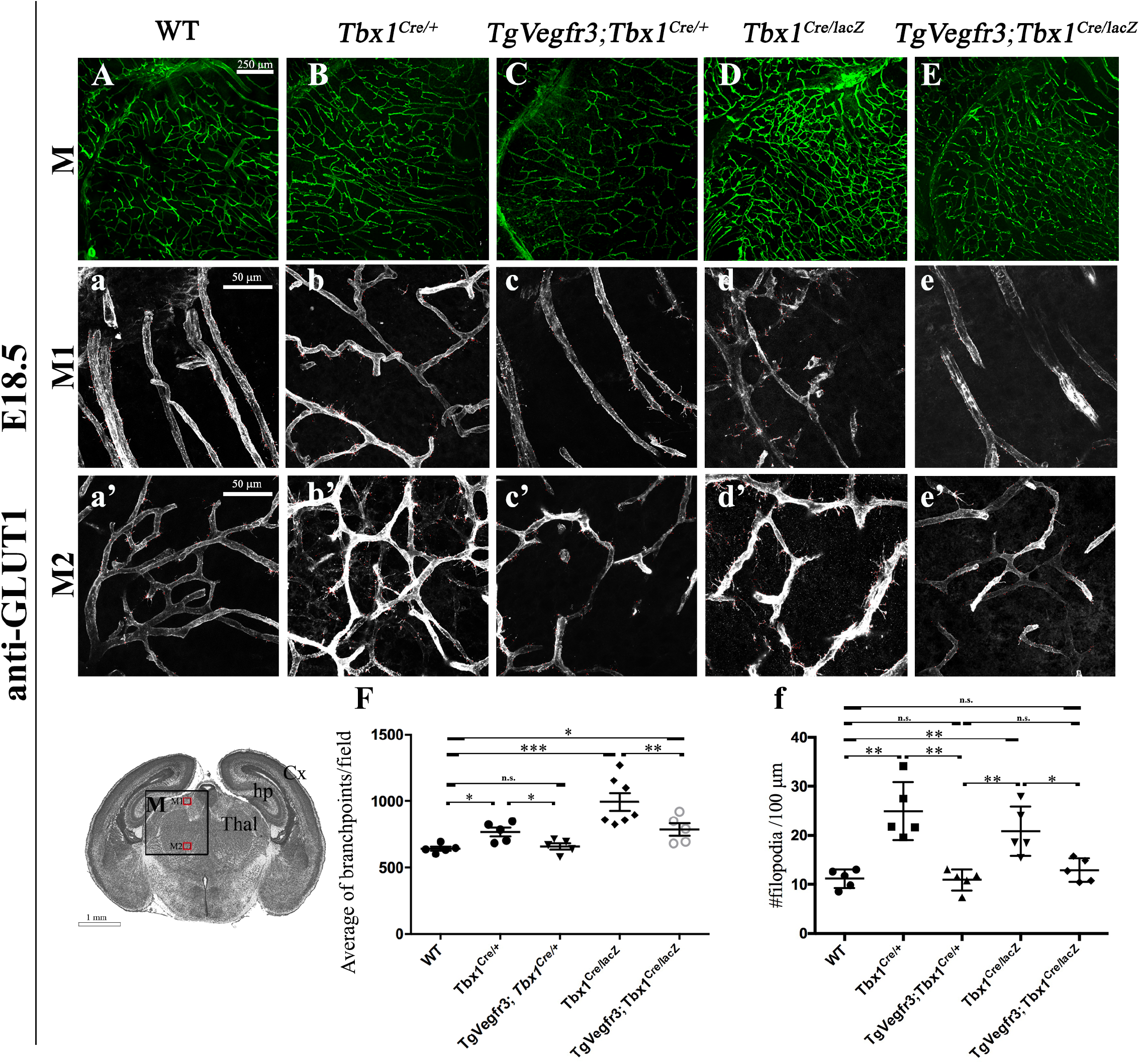
Vegfr3 over-expression rescues brain vessel abnormalities in Tbx1 mutants. (A - E) Representative coronal brain sections (medial) of embryos at E18.5 immunostained for GLUT1. The cartoon indicates the position of the counting boxes (M1, M2), shown at high magnification in panels M1 (a - e) and M2 (a’ - e’), that were used to quantify vessel branchpoint (F) and filopodial (f) density in embryos with the indicated genotypes. *** *P* value <0.001, ** *P* value <0.01, * *P* value <0.05. Error bars ± SD. Abbreviations: Cx, cortex, hp, hippocampus, Thal, thalamus.

In conclusion, the data from our loss and gain of function genetic experiments are consistent with a *Tbx1* > *Vegfr3* gene cascade that establishes the correct brain vessel density in the developing mouse brain. As full rescue of the brain vascular anomalies caused be *Tbx1* mutation was achieved by activating *Vegfr3* exclusively in *Tbx1*-expressing ECs, we propose that critical *Tbx1* functions in brain angiogenesis are cell-autonomous.

## DISCUSSION

*Tbx1* positively regulates three genes that suppress angiogenesis, *Vegfr3, Dll4* and *Unc5b* (Cioffi et al., 2014); the inactivation of any one of which leads to increased brain vessel density, similar to that observed in *Tbx1* mutants (Lu et al., 2004; Suchting et al., 2007; Tammela et al., 2008). We have shown that a JAG1 peptide only partially rescued microtubule hyperbranching in *Tbx1*-depleted cultured HUVECs (Cioffi et al., 2014), suggesting that Notch-independent pathways contribute to the pathological process. Here we investigated the TBX1-VEGF-C/VEGFR3 axis.

Because TBX1 can regulate *Vegfr3* expression at a transcriptional level (Chen et al., 2010), we hypothesize that this is the mechanistic basis of the observed interactions, but is this enough? There are several points to consider, i) the increased brain vessel density in *Tbx1*^*-/-*^ embryos is not severe, thus TBX1 appears to have only a limited, modulatory/suppressor function over vessel branching, ii) *Tbx1* positively regulates at least three genes that suppress angiogenesis, *Vegfr3, Dll4* and *Unc5b* (Cioffi et al., 2014); the inactivation of any one of which leads to increased brain vessel density, similar to that observed in *Tbx1* mutants (Lu et al., 2004; Suchting et al., 2007; Tammela et al., 2011), iii) a JAG1 peptide only partially rescued microtubule hyperbranching in cultured *Tbx1*-depleted HUVECs (Cioffi et al., 2014), suggesting that Notch-independent pathways are involved (hence investigating the VEGF-C-VEGFR3 pathway). Thus, while it is possible that the loss of *Tbx1* perturbs a genetic circuit modulating vessel branching through multiple mechanisms, our data demonstrate that enhanced expression of *Vegfr3* is sufficient to rebalance or override it and re-establish normal branching. Thus, we conclude that VEGFR3 is a critical target of *Tbx1* in brain microvessels.

We have gained insights into the timing of the critical *Tbx1* - *Vegfr3* interaction. We found VEGFR3 to be expressed in most brain vessels between E13.5 and E18.5. Thus, TBX1 and VEGFR3 co-localize in a subpopulation of brain ECs from the timepoint in which the brain vessel hyperbranching phenotype was first identifiable in *Tbx1* mutant embryos, namely at E15.5, suggesting that the co-presence of the two proteins may be part of the pathogenetic mechanism. The time course analysis of *Tbx1* cell fate mapping performed here showed that at E10.5, isolated GFP+ cells (*Tbx1*-expressing and their descendents) were within hindbrain vessels and in the PNVP, but we did not observe entire blood vessels that were GFP+. The latter would be expected if *Tbx1* were activated in EC progenitors. After E11.5, the number of GFP+ cells increased progressively in more rostral and dorsal brain regions, as illustrated in the cartoon in Fig. 1. Thus, the distribution of *Tbx1*-fated cells likely reflects a wave of activation that expands in different regions of the growing brain vascular network, rather than the deployment of *Tbx1*-expressing EC progenitors and their progeny. The fact that the brain vascular phenotype was only detectable after E13.5, is consistent with the interpretation that critical TBX1 functions are in differentiated ECs and not in EC progenitors, in contrast, for example to the role of TBX1 in cardiac lymphatic vessels (Martucciello et al., 2020).

There are no published studies of VEGFR3 function specifically in brain vessels. Some studies have suggested that VEGFR3 does not play a primary role in angiogenesis, and that the phenotypic consequences of its inactivation upon this process are due to modulation of VEGFR2. For example, targeted inactivation of the tyrosine kinase or the ligand-binding domain of VEGFR3 did not affect angiogenesis in mouse embryos or yolk sacs (Zhang et al., 2010). As VEGFR3 and VEGFR2 form heterodimers, inactivation of the *Vegfr3* gene might affect signalling through the VEGFR2 receptor. Our study does not exclude this possibility as inactivation of *Tbx1* in cultured HUVECs and in brain ECs isolated from *Tbx1* mutants led to increased expression of *Vegfr2* (Cioffi et al., 2014). Another intriguing possibility is that the bi-modal functions of VEGFR3 in ECs (pro- or anti-angiogenic) depend upon local levels of Notch (Benedito et al., 2012). Several studies have contributed to the definition of an autoregulatory loop in which VEGF-C-VEGFR3 signalling activates Dll4-Notch, which promotes the conversion of endothelial tip cells into stalk cells, and subsequently suppression of the VEGF receptors, including VEGFR3 (Benedito et al., 2012; Jakobsson et al., 2009; Tammela et al., 2008; Tammela et al., 2011), thereby preventing excessive angiogenesis. We have proposed that TBX1 operates upstream of this autoregulatory loop (Cioffi et al., 2014), because genetic inactivation of *Vegfr3* and *Dll4*, which are regulated by TBX1, and of *Tbx1* itself, all result in brain vessel hyperbranching (Cioffi et al., 2014; Suchting et al., 2007; Tammela et al., 2011), due presumably to the dominance of tip cells over stalk cells. The results of our current study sustain this hypothesis, because reduced expression of *Vegfr3* enhances the *Tbx1* mutant phenotype, while increased expression suppresses it. In addition, *Vegfr3* and *Tbx1* are co-expressed in tip cells, consistent with an interaction that represses vessel branching.

## ACKNOWLEDGEMENTS

We are grateful for the invaluable support provided by the Integrated Microscopy Core and the Animal Facility at the Institute of Genetic and Biophysics “ABT”/CNR, Naples and to Drs. Claudia Angelini and Annamaria Carissimo of the Istituto Applicazioni del Calcolo, CNR, Naples, for assistance with the statistical analysis. *Vegfr3*^*flox/+*^ mice were generously provided by Dr Kari Alitalo.

## SUPPLEMENTARY FIGURES

*Supplementary Figure S1. Distribution of Tbx1-fated cells*

(A - F) Tbx1-fated cells in rostral (R), medial (M) and caudal (C) coronal brain sections of *Tbx1*^*Cre/+;*^ *Rosa*^*mTmG*^ embryos at E18.5 immunostained with antibodies to GFP (green) and GLUT1 (red). Boxed areas in A, C, E, are enlarged in the adjacent panels. (G - L) Co-localization of GLUT1 and VEGFR3 in brain vessels in rostral (R), medial (M) and caudal (C) coronal brain sections of *Tbx1*^*Cre/+;*^ *Rosa*^*mTmG*^ embryos at E13.5. Boxed areas in G, I, K are enlarged in the adjacent panels. (M - R) Expression of GFP and VEGFR3 in brain vessels in rostral (R), medial (M) and caudal (C) coronal brain sections of *Tbx1*^*Cre/+;*^ *Rosa*^*mTmG*^ embryos at E18.5. Boxed areas in M, O, Q are enlarged in the adjacent panels. Abbreviations: Cx, cortex, GE, ganglionic eminences, Thal, thalamus, Hyp, hypothalamus, Pn pons, sc, superior colliculus.

*Supplementary Figure S2. Tbx1 and Vegfr3 interact genetically to regulate brain vessel and filopodial density*.

Representative coronal brain sections, in rostral (A - E) and caudal (F - J) positions of E18.5 embryos immunostained for GLUT1. The cartoon of a rostral brain section (R) indicates the position of the counting boxes R1 and R2, which are shown at high magnification in panels R1, a - e and R2, a’ - e’. The cartoon showing a caudal brain section (C) indicates the position of the counting boxes C1 and C2, shown at high magnification in panels C1, a - e and C2, a’ - e’. Abbreviations: Cx, cortex, GE, ganglionic eminences, sc, superior colliculus, Pn, pons.

*Supplementary Figure S3. Vegfr3 over-expression rescues brain vessel abnormalities in Tbx1 mutants*.

Representative coronal brain sections, in rostral (A - E) and caudal (F - J) positions, of embryos at E18.5 immunostained for GLUT1. The cartoon showing a rostral brain section (R) indicates the position of the counting boxes R1 and R2, which are shown at high magnification in panels R1, a - e and R2, a’ - e’. The cartoon showing a caudal brain section (C) indicates the position of the counting boxes C1 and C2, shown at high magnification in panels C1, a - e and C2, a’ - e’. Abbreviations: Cx, cortex, GE, ganglionic eminences, sc, superior colliculus, Pn, pons.

## REFERENCES

Benedito, R., Rocha, S. F., Woeste, M., Zamykal, M., Radtke, F., Casanovas, O., Duarte, A., Pytowski, B. and Adams, R. H. (2012). Notch-dependent VEGFR3 upregulation allows angiogenesis without VEGF-VEGFR2 signalling. Nature 484, 110–114.

Chen, L., Mupo, A., Huynh, T., Cioffi, S., Woods, M., Jin, C., McKeehan, W., Thompson-Snipes, L., Baldini, A. and Illingworth, E. (2010). Tbx1 regulates Vegfr3 and is required for lymphatic vessel development. J. Cell Biol. 189, 417–424.

Cioffi, S., Martucciello, S., Fulcoli, F. G., Bilio, M., Ferrentino, R., Nusco, E. and Illingworth, E. (2014). Tbx1 regulates brain vascularization. Hum. Mol. Genet. 23, 78–89.

Huynh, T., Chen, L., Terrell, P. and Baldini, A. (2007). A fate map of Tbx1 expressing cells reveals heterogeneity in the second cardiac field. Genesis 45, 470–475.

Jakobsson, L., Bentley, K. and Gerhardt, H. (2009). VEGFRs and Notch: a dynamic collaboration in vascular patterning. Biochem Soc Trans 37, 1233–1236.

Karkkainen, M. J., Ferrell, R. E., Lawrence, E. C., Kimak, M. A., Levinson, K. L., McTigue, M. A., Alitalo, K. and Finegold, D. N. (2000). Missense mutations interfere with VEGFR-3 signalling in primary lymphoedema. Nat. Genet. 25, 153–159.

Karkkainen, M. J., Saaristo, A., Jussila, L., Karila, K. A., Lawrence, E. C., Pajusola, K., Bueler, H., Eichmann, A., Kauppinen, R., Kettunen, M. I., et al. (2001). A model for gene therapy of human hereditary lymphedema. Proc. Natl. Acad. Sci. U.S.A. 98, 12677–12682.

Lindsay, E. A., Vitelli, F., Su, H., Morishima, M., Huynh, T., Pramparo, T., Jurecic, V., Ogunrinu, G., Sutherland, H. F., Scambler, P. J., et al. (2001). Tbx1 haploinsufficieny in the DiGeorge syndrome region causes aortic arch defects in mice. Nature 410, 97–101.

Lu, X., Le Noble, F., Yuan, L., Jiang, Q., De Lafarge, B., Sugiyama, D., Bréant, C., Claes, F., De Smet, F., Thomas, J.-L., et al. (2004). The netrin receptor UNC5B mediates guidance events controlling morphogenesis of the vascular system. Nature 432, 179–186.

Mäkinen, T., Jussila, L., Veikkola, T., Karpanen, T., Kettunen, M. I., Pulkkanen, K. J., Kauppinen, R., Jackson, D. G., Kubo, H., Nishikawa, S., et al. (2001). Inhibition of lymphangiogenesis with resulting lymphedema in transgenic mice expressing soluble VEGF receptor-3. Nat. Med. 7, 199–205.

Martucciello, S., Turturo, M. G., Bilio, M., Cioffi, S., Chen, L., Baldini, A. and Illingworth, E. (2020). A dual role for Tbx1 in cardiac lymphangiogenesis through genetic interaction with Vegfr3. FASEB J 34, 15062–15079.

Muzumdar, M. D., Tasic, B., Miyamichi, K., Li, L. and Luo, L. (2007). A global double-fluorescent Cre reporter mouse. Genesis 45, 593–605.

Ogata, T., Niihori, T., Tanaka, N., Kawai, M., Nagashima, T., Funayama, R., Nakayama, K., Nakashima, S., Kato, F., Fukami, M., et al. (2014). TBX1 mutation identified by exome sequencing in a Japanese family with 22q11.2 deletion syndrome-like craniofacial features and hypocalcemia. PLoS ONE 9, e91598.

Paxinos G, and Franklin K. (2012). Mouse brain in stereotaxic coordinates. New York.

Paylor, R., Glaser, B., Mupo, A., Ataliotis, P., Spencer, C., Sobotka, A., Sparks, C., Choi, C.-H., Oghalai, J., Curran, S., et al. (2006). Tbx1 haploinsufficiency is linked to behavioral disorders in mice and humans: implications for 22q11 deletion syndrome. Proc. Natl. Acad. Sci. U.S.A. 103, 7729–7734.

Suchting, S., Freitas, C., le Noble, F., Benedito, R., Bréant, C., Duarte, A. and Eichmann, A. (2007). The Notch ligand Delta-like 4 negatively regulates endothelial tip cell formation and vessel branching. Proc. Natl. Acad. Sci. U.S.A. 104, 3225–3230.

Tammela, T., Zarkada, G., Wallgard, E., Murtomäki, A., Suchting, S., Wirzenius, M., Waltari, M., Hellström, M., Schomber, T., Peltonen, R., et al. (2008). Blocking VEGFR-3 suppresses angiogenic sprouting and vascular network formation. Nature 454, 656–660.

Tammela, T., Zarkada, G., Nurmi, H., Jakobsson, L., Heinolainen, K., Tvorogov, D., Zheng, W., Franco, C. A., Murtomäki, A., Aranda, E., et al. (2011). VEGFR-3 controls tip to stalk conversion at vessel fusion sites by reinforcing Notch signalling. Nat. Cell Biol. 13, 1202–1213.

Torres-Juan, L., Rosell, J., Morla, M., Vidal-Pou, C., García-Algas, F., de la Fuente, M.-A., Juan, M., Tubau, A., Bachiller, D., Bernues, M., et al. (2007). Mutations in TBX1 genocopy the 22q11.2 deletion and duplication syndromes: a new susceptibility factor for mental retardation. Eur. J. Hum. Genet. 15, 658–663.

Watanabe, C., Matsushita, J., Azami, T., Tsukiyama-Fujii, S., Tsukiyama, T., Mizuno, S., Takahashi, S. and Ema, M. (2019). Generating Vegfr3 reporter transgenic mouse expressing membrane-tagged Venus for visualization of VEGFR3 expression in vascular and lymphatic endothelial cells. PLoS ONE 14, e0210060.

Yagi, H., Furutani, Y., Hamada, H., Sasaki, T., Asakawa, S., Minoshima, S., Ichida, F., Joo, K., Kimura, M., Imamura, S., et al. (2003). Role of TBX1 in human del22q11.2 syndrome. Lancet 362, 1366–1373.

Zarkada, G., Heinolainen, K., Makinen, T., Kubota, Y. and Alitalo, K. (2015). VEGFR3 does not sustain retinal angiogenesis without VEGFR2. Proc. Natl. Acad. Sci. U.S.A. 112, 761–766.

Zhang, L., Zhou, F., Han, W., Shen, B., Luo, J., Shibuya, M. and He, Y. (2010). VEGFR-3 ligand-binding and kinase activity are required for lymphangiogenesis but not for angiogenesis. Cell Res 20, 1319–1331.

